# High-throughput Automated Single Cell Imaging Analysis Reveals Dynamics Of Glioblastoma Stem Cell Population During State Transition

**DOI:** 10.1101/381715

**Authors:** Anastasia P. Chumakova, Masahiro Hitomi, Erik P. Sulman, Justin D. Lathia

## Abstract

Cancer stem cells (CSCs) are a heterogeneous and dynamic population that stands at the top of tumor cellular hierarchy and is responsible for maintenance of the tumor microenvironment. As methods of CSC isolation and functional interrogation advance, there is a need for a reliable and accessible quantitative approach to assess heterogeneity and state transition dynamics in CSCs. We developed a High-throughput Automated Single Cell Imaging Analysis (HASCIA) approach for quantitative assessment of protein expression with single cell resolution and applied the method to investigate spatiotemporal factors that influence CSC state transition using glioblastoma (GBM) CSC as a model system. We were able to validate the quantitative nature of this approach through comparison of the protein expression levels determined by HASCIA to those determined by immunoblotting. A virtue of HASCIA was exemplified by detection of a subpopulation of SOX2-low cells, which expanded in fraction size during state transition. HASCIA also revealed that CSCs were committed to loose stem cell state at an earlier time point than the average SOX2 level decreased. Functional assessment of stem cell frequency in combination with quantification of SOX2 expression by HASCIA defined a stable cut-off of SOX2 expression level for stem cell state. We also developed an approach to assess local cell density and found that denser monolayer areas possess higher average levels of SOX2, higher cell diversity and a presence of a sub-population of slowly proliferating SOX2-low CSCs. HASCIA is an open source software that facilitates understanding the dynamics of heterogeneous cell population such as that of CSCs and their progeny. It is a powerful and easy-to-use image analysis and statistical analysis tool available at https://hascia.lerner.ccf.org.

## INTRODUCTION

Heterogeneity is a predominant feature of most biological systems (1,2,3) and emerging evidence supports the functional importance of cellular heterogeneity and noise for stability and plasticity in cell populations (4, 5, 6). Heterogeneity of response ensures robustness and stability of a population; phenotypic and genomic heterogeneity provides a substrate for adaptation and populational evolution. Complex tissue microenvironments are characterized by cellular hierarchies and uneven distribution of ligands, metabolites and energy sources, which in turn further increases heterogeneity of such systems. In cancers, intratumoral heterogeneity results in multiple cell subpopulations (7). Implications of such diversity include development of therapeutic resistance, tumor evolution and cellular population drift. Many cancer types have been shown to exhibit remarkable cellular heterogeneity including glioblastoma (GBM). This primary malignant brain tumor is notorious for development of therapeutic resistance, genetic and phenotypic diversity and drastic temporal cellular drift (8).

Cancer stem cells (CSCs) have been first proposed as a model for acquisition of intratumoral heterogeneity (9). To date there is substantial evidence to support that GBM CSCs are involved in establishment of a diverse tumor microenvironment (10-15). Recent advances in isolation, manipulation and interrogation of CSCs have made it possible to recapitulate CSC behavior and study these cells in vitro (16).

Emergence of single cell assessment approaches enabled evaluation of cellular diversity. Single cell DNA and RNA sequencing as well as flow cytometry and mass spectrometry have become widely used methods in the field of cancer research and have revolutionized our understanding of CSCs (17-20). However, most of the research efforts focus on static assessment of CSCs and underestimate spatial distribution of cell population. GBM CSCs are a dynamic system: these cells can transition in and out of the stem cell state and form intricate communication network within CSC population as well as with other cell types within the tumor (21-24). However, there is a lack of experimental approaches to assess spatiotemporal aspects of CSC biology.

Immunofluorescence staining allows for direct visualization of cellular antigens in vivo and in vitro. The method has remained a staple of antigen detection and phenotype assessment since 1940 (25). However, despite widely accessible digital image acquisition, immunofluorescence implementation in modern research remains mostly qualitative. A survey of Pubmed using *“Journal Article” [PT] AND 2017 [DP] AND “immunofluorescence” [TW]* request revealed that only 13 out of 50 papers studied (26%) used immunofluorescence quantification techniques to report study results. Most of the 13 papers we reviewed utilized manual data quantification. An average modern smartphone possesses 2.5 times more computing power than 1985 supercomputer (26), therefore computation speed and digital memory no longer pose limitations for most of the applications in modern science. In most areas of research the real limitation comes from lack of accessible, user-friendly software.

Large-scale programmatically-driven data analysis is being used progressively more in biology in areas such as genomics, transcriptomics and mass spectrometry (27-30). Programming languages and computing environments like R, Python and Matlab offer powerful tools for data processing and analysis (31-33). Scientific image analysis has become accessible through development of ImageJ - open source software by NIH (34, 35). ImageJ and its specific application for biology - Fiji (36) allows users to quantify a wide variety of parameters based on pixel intensity and image coordinates. The software offers tools and plugins for particle analysis and an easy-to-learn macro language for batch image processing. RStudio’s Shiny is a platform for development of user-friendly interfaces for R-based statistical inference and data visualization.

We utilized ImageJ macro and R languages, as well as the Shiny platform to create an accessible open source high-throughput automated single cell imaging analysis (HASCIA) approach to immunofluorescence quantification. We applied the method to investigate population dynamics during GBM CSC state transition and examined spatiotemporal factors that play a role in this process.

## MATERIALS AND METHODS

### CSC derivation and xenograft maintenance

Previously established patient-derived xenograft (PDX) model of GBM (sample MDA23) (37) was obtained via a material transfer agreement from The University of Texas MD Anderson Cancer Center, Houston, TX. GBM cells were maintained using subcutaneous xenografts as previously described (38, 39). All experiments utilizing mice were approved by the Institutional Animal Care and Use Committee (IACUC) of the Cleveland Clinic. PDX model was maintained using immune-deficient NOD.Cg-Prkdcscid ||2rgtm1Wj|/SzJ (NSG) mice (Jackson Laboratory, Bar Harbor, ME, USA) for maintenance of tumor heterogeneity. Six-week-old female mice were subcutaneously injected in the flank with 2*10^6 freshly dissociated PDX GBM cells. Animals were sacrificed when tumor volume exceeded 5% of the animal’s body weight. Xenografted tumors were dissected mechanically and using papain dissociation kit (Worthington Biochemical Corporation, Lakewood, NJ, USA), CD133+ cells were subsequently enriched using the CD133 Magnetic Bead Kit for Hematopoietic Cells (CD133/2; Miltenyi Biotech, San Diego, CA, USA).

GBM CSCs were propagated as a non-adherent sphere culture using Neurobasal complete medium (NBC) (Neurobasal medium (Life Technologies, Carlsbad, CA, USA) supplemented with B27 (Life Technologies), 1% penicillin/streptomycin (Life Technologies), 1 mM sodium pyruvate (Life Technologies), 2 mM L-glutamine (Life Technologies), 20 ng/mL EGF (R&D Systems, Minneapolis, MN, USA), and 20 ng/mL FGF-2 (R&D Systems) in humidified incubator with 5% CO2. Only low passage cells (<10) were used in all experiments to prevent cellular drift.

### CSC treatments

For immunofluorescence staining and immunoblotting experiments cells were cultured as adherent monolayer on plates or glass cover slips coated Geltrex (Thermo Fisher Scientific, Waltham, MA, USA). To demonstrate change in epidermal growth factor receptor (EGFR) activation cells were cultured in three different conditions for 3 days: 1) NBC - Neurobasal complete medium: supplemented with 20 ng/mL EGF, and 20 ng/mL FGF-2, B27, 1% penicillin/streptomycin, 1 mM sodium pyruvate, and 2 mM L-glutamine; 2) NBF - Neurobasal medium supplemented with 10% fetal bovine serum (FBS, Sigma Aldrich, St. Louis, MO, USA), B27,1% penicillin/streptomycin, 1 mM sodium pyruvate, and 2 mM L-glutamine; 3) DMEM+FBS - Dulbecco’s modified Eagle’s medium (DMEM) supplemented with 10% FBS and 1% penicillin/streptomycin. For state transition experiments NBC with 25 ng/mL of bone morphogenic protein 4 (BMP4, R&D Systems) was used for 1-5 days to induce CSC differentiation.

### Immunoblotting

Whole cell lysates were collected from adherent monolayer cell culture using 10% NP40 (Sigma Aldrich), 1mM EDTA, 150 mM NaCl, 10mM TrisCl pH 7.5 lysis buffer supplemented with protease and phosphatase inhibitor cocktails (Sigma Aldrich). GBM CSCs were analyzed by immunoblotting for expression of phospho-EGFR (Y1068, 1:1000, Cell Signaling, Danvers, MA, USA), SOX2 (1:500, R&D Systems) and phospho-STAT3 (1:1000, Y705, Cell Signaling). Anti-beta-Actin (1:5000, Santa Cruz Biotechnology, Dallas, TX, USA) was used for loading control.

### Limiting dilution assay

After exposure to one to five days of BMP4 cells were plated in a 96 well format with 24 wells of each dilution: 0.78, 1.56, 3.125, 6.25,12.5, 25, 50 and 100 cells/well. Number of wells positive for spheres was calculated 14 days later and stem cell frequencies were calculated using online tool available through the Walter and Eliza Hall Institute of Medical Research (http://bioinf.wehi.edu.au/software/elda/index.html) (40).

### Proliferation assay

CellTrace Far Red dye (Life Technologies) was used for tracing of cell proliferation activity within the five-day period. Adherent monolayer cells were incubated with 1:1000 dilution of the dye concentrate in Neurobasal medium for 20 min at 37 C. Following 3X wash with fresh Neurobasal medium cells were incubated in NBC medium for five days. Cells retaining the highest amount of dye represented the slowly dividing pool where as cells that divided multiple times had diluted the dye and therefore presented lower fluorescence intensity in far red channel.

### HASCIA method description

#### Phase I: Immunofluorescence staining

To enable high-throughput parallel analysis of multiple conditions and up to four technical replicates we cultured cells in a 24-well format with glass cover-slips (Electron Microscopy Sciences, Hatford, PA, USA) on the bottom of the wells. To ensure stem cell phenotype is preserved during propagation as an adherent monolayer we coated the glass cover-slips with laminin-rich Geltrex solution (Life Technologies). Cells were fixed with a 4% solution of paraformaldehyde in phosphate buffered saline. After blocking and permeabilization with 2% donkey serum (EMD Millipore) and 0.1% triton (Sigma) we incubated the cells with 150 μL of primary and subsequently 500 μL of secondary antibody solutions directly in the 24-well plates, washing 3X with 1000 μL of PBS in between. Primary antibodies used: phospho-EGFR (Y1068, Cell Signaling), SOX2 (R&D Systems) and phospho-STAT3 (Y705, Cell Signaling). Secondary antibodies (Jackson ImmunoResearch): DL-488-conjugated donkey anti-mouse IgG, Cy3-conjugated donkey antimouse IgG, DL-649-conjugated donkey anti-rabbit IgG.

As the last step, the internal controls (DNA and in some cases - Actin) were stained by incubating the cells with Hoechst 33342 (Polyscience Inc, Warrington, PA, USA) and phalloidin-Alexa-594 (Invitrogen, Carlsbad, CA, USA) solutions in PBS. Cover-slips were picked up from the wells and mounted on slides using gelvatol mounting medium (PVA (Sigma), Glycerol (Sigma), Sodium Azide (Fisher), TrisCI ph 8.5, water).

#### Phase II: Digital imaging

Imaging was carried out using Leica DM5000B microscope equipped with a Leica DFC310 FX Digital Color Camera (Fig. 1A). Five to twenty fields of view were captured from every cover-slip in multiple channels.

**Figure 1.**
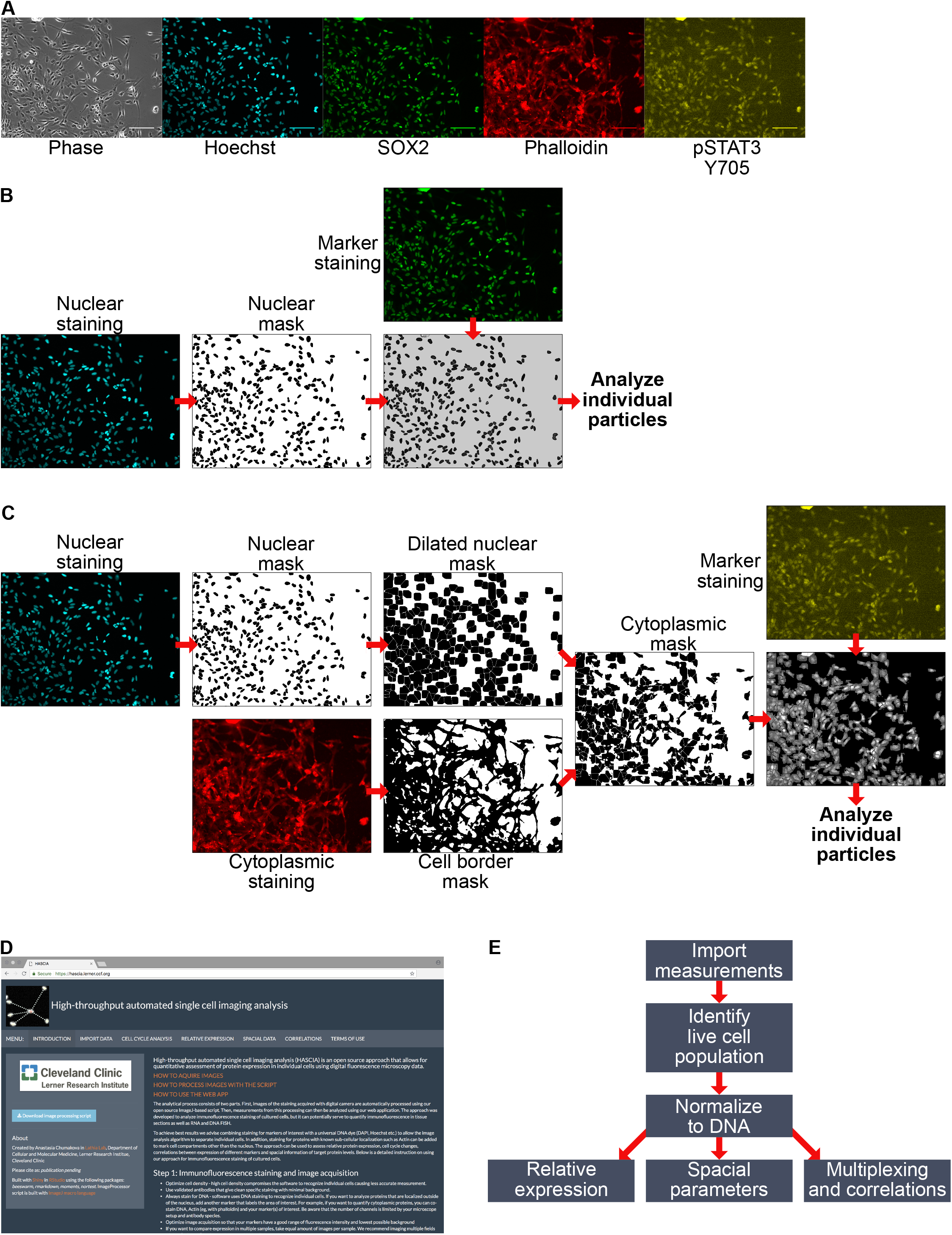
High-throughput automated single cell imaging analysis (HASCIA) method description. **A** - immunofluorescence images of an adherent GBM CSC monolayer, scale bar – 100 μm. Phase – phase contrast, Hoechst – DNA staining, SOX_2_ – stem cell marker, Phalloidin – b-Actin staining, PSTAT_3_ Y705 - phosphorylated Y705 of STAT3 staining. **B** – Algorithm for nuclear staining analysis. Nuclear staining images are used to create nuclear masks and separating particles. The mask is further used for delineation of nuclear region in nuclear marker staining. Individual particles are subsequently analyzed using Particle Analysis feature of ImageJ. **C** – Algorithm for cytoplasmic staining analysis. Nuclear mask is created as in B with subsequent dilation to fill perinuclear area. Cytoplasmic standard staining (in this case ’ Actin, stained by Phalloidin-Alexa594) is used to create cell border outlines. Combined dilated nuclear mask and cell border mask yield cytoplasmic mask that is then used to delineate individual cellular cytoplasm on images of marker staining. **D** – screen shot of https://hascia.lerner.ccf.org. **E** – Algorithm for HASCIA web-application data processing. After image processing with ImageJ scrip, measurements are imported into the app, based on DNA intensity profiles live cell population for every specimen is outlined, expression in specimens is calculated by applying normalization of marker intensity to DNA intensity in each individual particle. The normalized data can further be used to compare expression between specimens, analyze local cell density and perform multiplexing to investigate correlations between parameters.

#### Phase III: Image processing

An ImageJ-based image processing script was developed to enable analysis of digital images.

Script was created using ImageJ macro language (35, 41) and under Fiji version 1.0 (36), additional plugins used: CLAHE (42). The flow of the script is shown below and in Fig. 1B-C, code can be found by downloading the script file from https://hascia.lerner.ccf.org.

1. Images from each condition and staining are grouped into stacks to enable batch analysis;
2. Images are then normalized using shading images to cancel out any illumination, optics, camera and /or shutter imperfections;
3. Based on the Hoechst DNA staining (nuclear internal control) a mask is created that delineates nuclear region and separates cells into individual particles using watershed functionality of ImageJ (Fig. 1B).
4. If the cytoplasmic staining needs to be analyzed, delineation of cell borders is performed using dilated nuclear mask and a mask of a cytoplasmic standard (phalloidin). (Fig. 1C)
5. The mask(s) is then applied to the staining of interest to measure fluorescence intensity in individual cells. The measurement is carried out using Analyze particles command optimized for automatic batch analysis of multiple image stacks.

The measurements obtained from image analysis include an image Label, Raw Integreted Density (RawlntDen) - a measure of intensity of fluorescence in each particle, Xand Y position of each particle on the image, Feret diameter - a maximum particle caliper, and many additional parameters that can be used further for data analysis.

#### Phase IV: Data analysis

For analysis of single cell measurements we developed a web-application using R and Shiny platform and packages *{beeswarm}, {rmarkdown}, {moments}, {nortest}, {shiny}* (Fig. 1D-E). Public version of all the software is available at https://hascia.lerner.ccf.org

Measurements from each marker as well as internal control created by image processing script are imported into the platform, organized and then processed to analyze the following:

1. Cell cycle changes. DNA staining intensity is plotted as a histogram or as a Kernel density plot. The web-app attempts to automatically identify G1 population by identifying the highest peak. G2 peak is identified by multiplying G1 peak x value by 2 (*G_1_x__* * *2*). Boundaries of live cell population are then calculated with the following formula and applied to the plot:

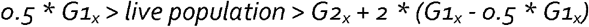 Each graph can be manipulated to manually select G_1_ peak or manually outline viable cell population and exclude dead cells and cell aggregates from further analysis.
2. Relative expression of stained markers. Using RawIntDen as a measure of fluorescence intensity, level of expression in different tested conditions can be compared. There are also two possible approaches for normalization of fluorescence intensity to internal control (i.e. DNA). The DNA intensity can be normalized to maximum of the selected live cell population and the marker intensity is divided by this normalized DNA value. Alternatively, marker intensity can be directly divided by DNA intensity. Both approaches enable more accurate per particle assessment of expression. Statistical analysis can be performed in this section: descriptive statistics include calculation of diversity coefficients, tests for normality, % outliers. Relative expression between conditions can be compared using t-test, pairwise t-test, Mann-Witney-Wilcoxon test and Kruskal-Wallis test.
3. Spatial distribution of cells. Image Label is specified to identify cells located on the same image. Then, based on X and Y coordinates position of the center of each particle is identified. Feret’s diameter is used as a measure of particle’s caliper and is used to obtain mean particle diameter. Two parameters can be calculated by our web-application: mean/median distance to top n number of closest neighbors (MDn) and number of particles within a certain radius of each cell (NWRn). Graphic representation of these parameters are outlined in Fig. 4A. Similarly to Relative expression section, comparative and descriptive statistical analysis can be performed on spatial parameters.
4. Correlations between multiple parameters including normalized and non-normalized marker expression, MDn and NWRn. Web application plots total population distribution and calculates Pearson’s correlation. A subset of the total population can be outlined and plotted independently.

### Further statistical analysis

For calculation and graphical presentation of diversity parameters we processed data using R, RStudio interface and *{base}* R graphics. To characterize differences in cell populations we used the following parameters:

1. Tsallis entropy (43, 44) using *Tsallis()* function from *{entropart}* R package;
2. Renyi entropy (45) using *Renyi.z()* function from {*EntropyEstimation}* R package;
3. Percent outliers (46) by determining Q1 (25%) and Q3 (75%) by *quantile()* function from *{stats}* and calculating interquartile range (IQR):

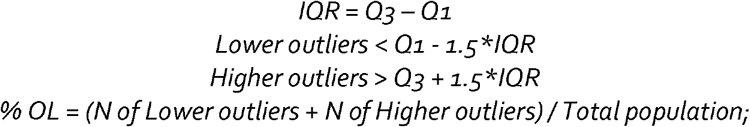
4. Kolmogorov-Smirnov test (47) was performed by comparing sample distribution with a normal distribution with the same average and standard deviation using functions *rnorm()* and *ks.test()* from *{stats}* R package;
5. Anderson-Darling test (48) was performed using *adtest()* from *{nortest}* R package.

## RESULTS

### HASCIA method validation and assessment of GBM CSC population heterogeneity

As a model of heterogeneous cell population we used PDX GBM CSCs. To validate the accuracy of assessing protein expression using HASCIA we first compared cumulative protein expression levels determined by immunoblotting with mean immunofluorescence staining intensities measured by HASCIA. For this purpose we chose a signaling pathway that is well-described and highly active in GBM CSC population: EGFR activation. To stimulate EGFR activation we stimulated with CSC NBC medium, which contains EGF. We used differentiation medium (DMEM+FBS) as well as Neurobasal medium without EGF and FGF but with addition of FBS (NBF) as a control (Fig.2A). Quantification of immunofluorescence staining for Tyr 1068 phosphorylated EGFR (pEGFR) using HASCIA revealed increased activation of the receptor in CSC propagating NBC medium that contains EGF. Interestingly, insulin-containing (through B27 supplement) NBF medium was able to partially increase pEGFR levels compared to DMEM+FBS (without B27) (Fig. 2A). The same EGFR activation pattern detected by immunoblotting in the parallel samples (Fig. 2B) validated that HASCIA can quantitatively determine the average protein expression of the population. Beyond the mean expression levels of pEGFP of the cultures, HASCIA revealed heterogeneity of single cell pEGFR/DNA level. We utilized statistical approaches described previously (46) to assess and compare heterogeneity of pEGFR expression in GBM CSC populations (Table 1). We compared diversity by calculating Tsallis and Renyi entropies and percent outliers in the three populations. Diversity was the highest in NBF differentiation inducing condition indicating that this medium was able to sustain wider spectrum of cell state. We were also able to assess normality of the expression distributions by Kolmogorov-Smirnov and Anderson-Darling tests. From these experiments it became apparent that when subjected to different media conditions response of the GBM CSCs population is heterogeneous rather than binary. Population responds not only by shifting average expression but also by increasing or decreasing diversity and number of outliers.

**Table 1.** Diversity and normality of pEGFR expression distribution in GBM CSCs

|  | DMEM+FBS | NBC | NBF |
| --- | --- | --- | --- |
| Tsallis entropy | 1.4969 | 1.9069 | 1.9276 |
| Renyi entropy | 1.2773 | 1.7189 | 1.7413 |
| % Outliers | 2.7701 | 2.4705 | 2.7331 |
| Kolmogorov-Smirnov index | 0.0886 | 0.0634 | 0.0932 |
| Kolmogorov-Smirnov p-value | 0.0069 | 0.0015 | 0.0090 |
| Anderson-Darling index | 19.0460 | 9.3235 | 11.7658 |
| Anderson-Darling p-value | 3.70E-24 | 1.37E-22 | 3.70E-24 |

**Figure 2.**
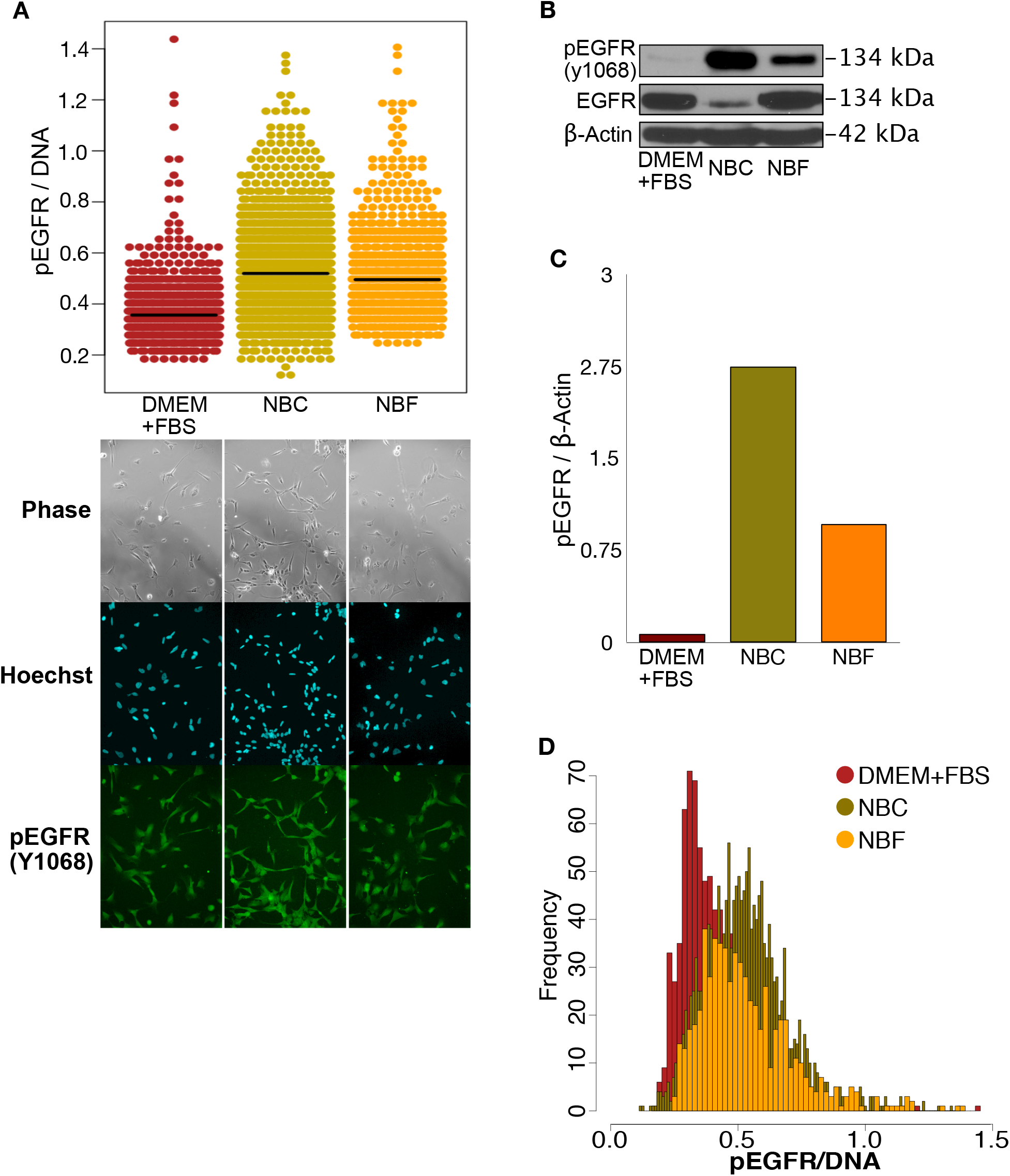
HASCIA method validation and assessment of GBM CSC population heterogeneity. **A** – Immunofluorescence staining and quantification using HASCIA. Adherent CSC monolayer was treated for 3 days with DMEM+FBS, EGF-containing NBC or with NBF medium. The cells were then fixed and stained for phosphorylated Y1068 EGFR (green) and using Hoechst (blue) as a nuclear standard. Quantification with HASCIA revealed significant differences in pEGFR expression between the three groups. **B** – In parallel, after similar treatment an immunoblotting analysis was performed with whole cell lysates. **C** – Densitometry analysis of the immunoblotting revealed similar trend of pEGFR changes with different media. **D** – More detailed analysis of expression distribution measured by HASCIA revealed differences in diversity and distribution shape in addition to changes in average pEGFR expression.

### Direct transition out of stem cell state

We next applied our method to investigate how GBM CSC population responds to a differentiation stimulus over time. We first determined SOX_2_ and PSTAT_3_ expression during a differentiation time courses as these markers have been described as predictors of sternness in CSC_s_ (49, 50) (Fig. 3A). In a direct time course GBM CSCs were subjected to bone morphogenic protein _4_ (BMP_4_) for one to five days and the cells were fixed and stained (Fig. 3B-C). Upon staining for SOX_2_ we observed a stable average SOX_2_ expression for the first three days and then a significant decline on days 4 and 5 (Fig. 3B). Interestingly, plotting SOX_2_ expression against PSTAT_3_ expression not only confirmed a strong correlation between the two markers, but also revealed a sub-population of cells that expressed two markers at distinctively low levels separating this sub-population of cells from the majority. We measured the percentage of the SOX_2_-low/pSTAT_3_-low sub-population and plotted it against the time of treatment (Fig. 3C). An increase in SOX_2_-low population was observed beginning day 2 of exposure to differentiation medium, suggesting a possible state transition in the population during this time. The expansion of the SOX_2_-low cell population preceded average SOX_2_ expression change by two days, which may indicate that state transition happens earlier than the cumulative marker expression change in the total population becomes apparent. Alternatively these data may indicate a sub-population with different kinetics of state transition.

**Figure 3.**
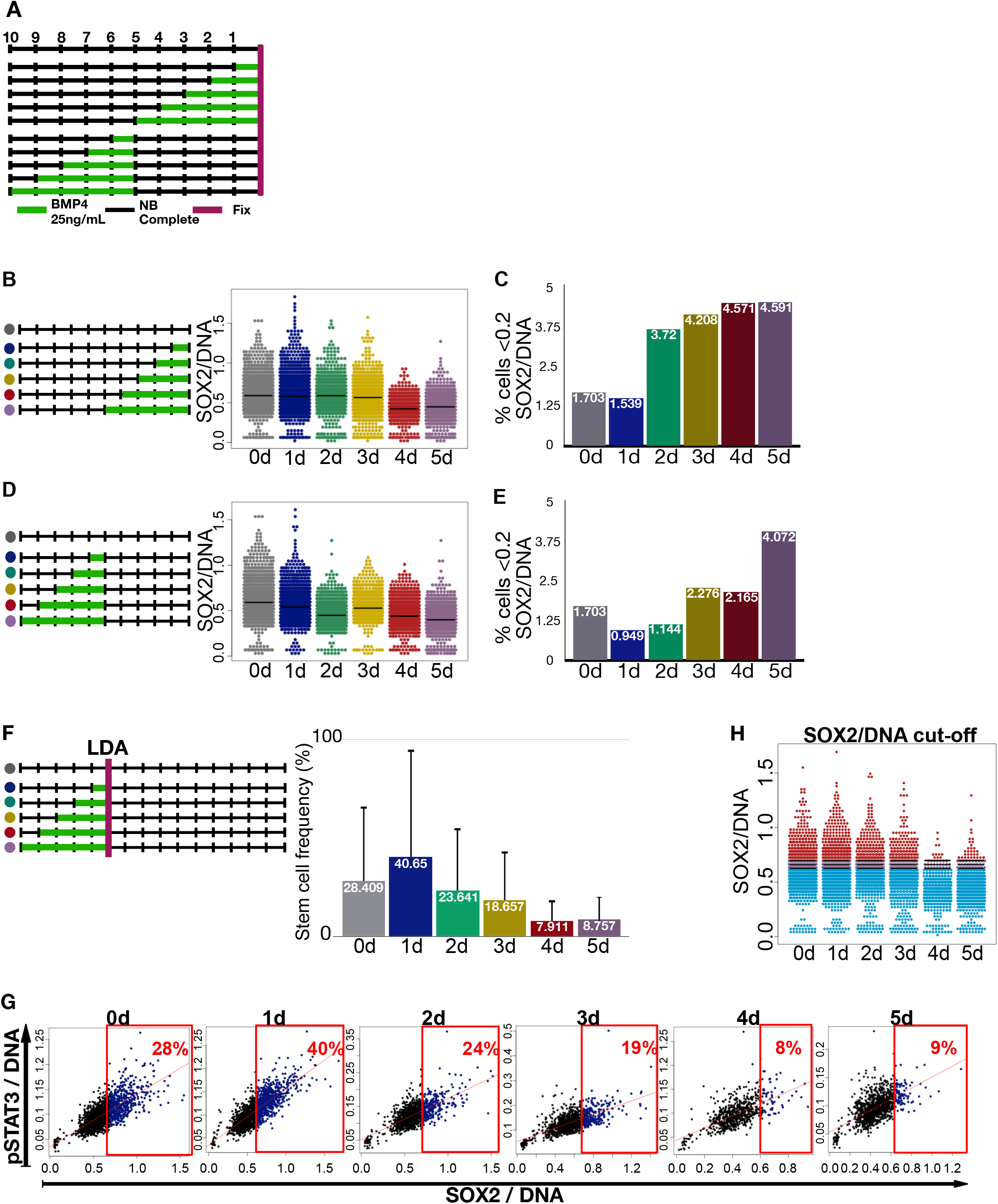
GBM CSC state transition and identification of SOX2 expression level in stem cell state. **A** – Experimental design for state transition time courses. GBM CSCs were cultured for total of 10 days in a 24-well plate. For differentiation induction experiments cells were treated with NBC + BMP_4_ for the last 0 – 5 days and then fixed and stained. For reversal of differentiation experiment cells were treated for 0 - 5 d of NBC + BMP_4_ and then returned to CSC-propagating NBC medium for 5 days before fixing and staining the cells. **B** – Differentiation induction time course. Overall SOX_2_/DNA levels of the population showed significant decrease on days 4 and 5. **C** – Quantification of SOX_2_-low population (<0.2 SOX_2_/DNA) revealed increase of this subpopulation percentage on day 2. **D** – Pre-treatment and reversal time course showed significant decrease in SOX_2_/DNA even after 1 day of pre-treatment with BMP4. **E** – Stable increase in percentage of SOX_2_-low subpopulation (<0.2 SOX_2_/DNA) was observed only after 5 days of pre-treatment. **F** – Functional assessment of sphere forming capacity using LDA estimated stem cell frequency of cultures treated with BMP_4_ for varying durations. **G** – Multiplexing of SOX_2_/DNA levels with pSTAT_3_/DNA from the direct time course (in A) helped outline estimated % of true stem cells (in D) and this way cut-off values of SOX_2_/DNA were determined. **H** – For each culture with various duration of BMP4 treatment, top percentile, which correspond to sphere forming frequency **(panel F)**, of CSCs racked according to SOX_2_ expression level were high lighted with red color, and the rest of the population was high lighted with blue color. Average cut-off between cut-offs of day 0-5 +/- standard deviation (purple) was calculated. This cut-off value changed very little and independent of BMP4 exposure duration suggesting that the cells express SOX_2_ at higher than this level are capable of sphere formation. Thus, it divides each cell population into sphere-forming stem cells (red) and non-sphere forming cells (blue).

### State transition and recovery during reversal of differentiation

In attempt to identify a temporal point of no return for CSC state transition we subjected cells to BMP_4_ pretreatment for one to five days, after which we returned stem cell propagating medium (NBC) for five days (Fig. 3D-E). Analysis if SOX_2_ expression using HASCIA image processing revealed that even after one day of BMP_4_ treatment and subsequent recovery for five days average SOX_2_ level was significantly lower that in non-treated cells (Fig. 3D). The longer the period of pretreatment the lower the average SOX_2_ was observed. We also performed SOX_2_-low/PSTAT_3_-low sub-population quantification similar to the direct transition experiment (Fig. 3E). Surprisingly, only cells pretreated for five days exhibited drastic increase in SOX_2_-low population. These data together with the results of direct transition out of stem cell state may indicate that marker expression changes are delayed in time from the actual state transition. To test that we performed a functional assessment of stem cell frequency using limiting dilution assay (LDA) (Fig. 3F). Estimated frequency of stem cells gradually decreased after day 2 of BMP_4_ exposure, which coincided with SOX_2_-low population expansion during direct time course. Together data from the time-courses and functional assay may indicate the beginning of state transition at day 2 with overall populational change in marker expression took place around day 4 - 5.

### Identifying SOX2 expression level in a stem cell state

Since the time course and self-renewal (LDA) experiments were performed using parallel replicates from a common GBM CSC population, we were able to directly compare the SOX_2_ expression levels with functional analysis. Using multiplex functionality of the HASCIA web-app we plotted the cell populations from the time course data of direct state transition experiment. We then outlined the top SOX2 expressing cells of the percentile that corresponded to the stem cell frequency determined by LDA for each time point (Fig. 3G). As a result we identified a cut-off value of SOX_2_/DNA for each population. Interestingly, the cut-off did not change in populations exposed to BMP_4_ for different length of time (Fig. 3H). This finding strongly supports SOX_2_ as an accurate marker of stem cell state and suggests that during state transition CSC population looses SOX_2_-high stem cells either through selection or through modulation of expression.

### Role of cell density in stem cell diversity and state transition

Importance of cell-to-cell contact and interaction with extracellular matrix for GBM CSC propagation has been outlined in a several recent studies (51-53). HASCIA processing script measurements for each particle include coordinates of each nucleus centroid in the field of view as well as information about the diameter of the particle. We used these measurements to develop an automated quantification of two density parameters (Fig. 4A). We hypothesized that cell density influences stem cell state and state transition. First we created an uneven culture of GBM CSCs with areas of low and high cell density (Fig. 4B). We compared SOX_2_ expression with median distance to top ten neighboring cells (MD_10_) and number of cells within four nuclear radii (NWR_4_). In both cases we found a weak correlation between the two parameters. The majority of SOX_2_-high cells were located in high-density areas. Subsetting cells from high-density (MD10 < 50 - Fig. 4C) and low-density (MD10 > 50 - Fig. 4D) areas revealed a significant difference in diversity (Tsallis and Renyi entropies) of these two sub-populations. We then looked at proliferation of cells depending on density. Using CellTrace dye retention as a marker of slowly dividing cells we cultured GBM CSCs for five days. As expected, high-density areas of the monolayer contained slowly dividing cells (Fig. 4E). Interestingly, we observed SOX_2_-low population to be slower dividing. The results suggest that high-density areas are more diverse and contain both slowly dividing differentiated SOX_2_-low cells and rapidly dividing SOX_2_-high CSCs. Low density areas on the contrary have lower SOX_2_ expression diversity, slightly lower average SOX_2_ expression with rapidly proliferating cells.

**Figure 4.**
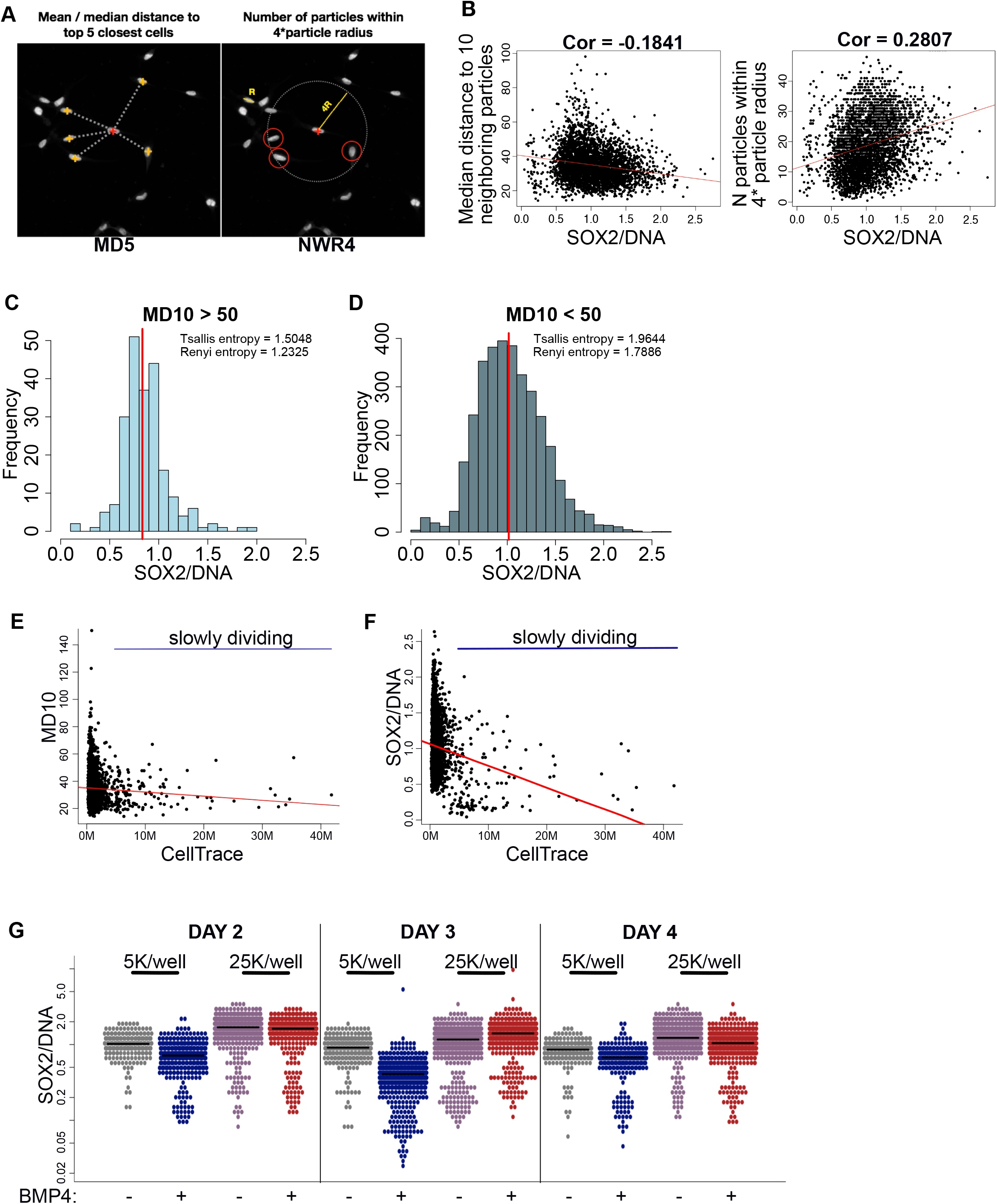
Cell density correlates with SOX2 level. **A** – Using particle coordinates HASCIA allows to quantify two spatial parameters: mean/median distance to top N neighboring particles (MDn) and number of particles within N * average particle radius (NWRn). **B** – SOX_2_/DNA was plotted against MD10 and NWR_4_, which revealed significant correlation (Pearson’s test) between these parameters. Highest SOX_2_/DNA was observed in high-density areas. **C** – Subset of CSC population in low local density (MD10<50) has a lower diversity (Tsallis and Renyi entropies) and slightly a lower average SOX_2_/DNA than **D** – CSCs in high local density area (MD10>50). **E** – 5 day proliferation assay revealed slowly dividing cells (high retention of CellTrace dye) in high-density areas. **F** – Slowly dividing population possesses lower SOX_2_/DNA level. **G** – State transition time course of cells plated at 5,000/well and 25,000/well revealed faster response in low-density culture and higher, more stable SOX_2_/DNA level in high-density culture.

Lastly, we looked at how local microenvironment affects state transition (Fig. 4G). Using BMP_4_ to facilitate differentiation we plated cells into a 24 well plate at two different densities - 5,000 cells/well and 25,000 cells/well - and measured SOX_2_ expression on day 2, 3 and 4. Consistent with findings in previous experiments low-density monolayer had lower average SOX_2_ expression and response to BMP4 was apparent on day two in this population. We found that high-density culture responded to differentiating condition two days later than low-density culture.

## DISCUSSION

For accurate assessment and modeling of heterogeneous cell population dynamics experimental approaches should allow for quantifiable results to be acquired. Qualitative approaches such as immunoblotting and classical qPCR assess cumulative or average genetic makeup or phenotype in a population based on assessment of total cell lysates. Furthermore, biological systems present with a significant level of noise, thus high-throughput analysis techniques are warranted for multiple replicates to be included. Single cell mass spectrometry and flow cytometry are methods that allow analyses with single cell resolution, but their implementation is often limited due to high cost of equipment and dedicated software and technical complexity that requires users to undergo specific training. Immunofluorescence staining is a well-established and relatively inexpensive approach, readily available in majority of cancer research laboratories. We present a solution for adapting this method to serve CSC research purposes with a single cell resolution.

Our open source software is easy to follow and does not require lengthy training. Our results are consistent with previously published papers (54-56) suggesting that quantitative immunofluorescence staining is reliably comparable to cumulative expression analysis by immunoblotting. It allows for better data normalization, normalization to DNA content, which cancels out the general increase in expression of many gene products due to cell cycle progression towards G2 cell cycle phase. Single particle resolution allows to assess heterogeneity of protein expression and it’s shift during response to biological stimuli in a cell population. Several software solutions allow automated image quantification however they all require development statistical analysis pipelines by the user (57-58). HASCIA yields statistically analyzed and graphically presented data, ready for publication. The software can potentially be optimized and used to analyze images of tissue sections and RNA and DNA FISH (59).

Heterogeneity of a cell population is often a result of a distinct minority sub-population being included in the total cell pool. Our results revealed a minor sub-population of GBM CSCs that is stable in a CSC propagating medium. Previous studies have suggested that multiple CSC subtypes may co-exist in the same tumor (60, 61). Observed increase in this minor SOX_2_-low sub-population followed BMP4-induced state transition suggests that these cells may be further on a differentiation spectrum.

Temporal aspects of commitment for stem state transition in CSCs are understudied. Our results suggest that even though phenotypical shift in a population can be observed following up to 4 days of continuous differentiation stimulus, partial irreversible commitment to differentiation can happen following just one day of stimulation with a significant decrease in SOX_2_ appearing even after recovery. These results underline the long lasting effects of microenvironment on CSCs and the extent of population plasticity in PDX GBM CSCs. SOX_2_ is an established progenitor cell program transcription factor in GBM CSCs (62, 63). Our results provide insight into a tight correlation between state transition and SOX_2_ expression. Narrow variation in identified SOX_2_ cut-off value defined by functional analysis supports direct influence of this protein expression on sphere and tumor forming capacity.

Cell density is closely dependent on cell adhesion and cell-to-cell communication. Our group and many other researchers have previously outlined the importance of adhesion and cell communication molecules in CSC maintenance and expansion (64-69). However in an experimental setting cell density is often overlooked and poorly controlled for across studies in CSC field. By developing a coordinate analysis of immunofluorescence images we were able to show that the local cell density impacted state transition dynamics, cellular diversity and cell proliferation. Optimal cell density for every particular GBM CSC specimen should be sought for reliable representation of GBM biology in vitro.

In conclusion, our results obtained via HASCIA underline the biological importance of CSC population heterogeneity. HASCIA can serve a useful and accessible tool to further interrogate heterogeneous and complex CSC behavior and is available through https://hascia.lerner.ccf.org

## ABBREVIATIONS

BMP4: bone morphogenic protein 4
CSC: cancer stem cell
EGFR: epidermal growth factor receptor
FBS: fetal bovine serum
GBM: glioblastoma
HASCIA: high-throughput automated single cell imaging analysis
IQR: interquartile range
LDA: limiting dilution assay
MDn: median distance to top N neighboring particles
NB: Neurobasal
NWRn: number of particles within N * average particle radius
OL: outliers
PDX: patient-derived xenograft

